# The right microbial stimulus can direct innate immune effector cells to specific organ sites to clear pathology

**DOI:** 10.1101/612598

**Authors:** Shirin Kalyan, Mark Bazett, Ho Pan Sham, Momir Bosiljcic, Beryl Luk, Salim Dhanji, Amanda M. Costa, Stephanie WY Wong, Mihai G. Netea, David W. Mullins, Hal Gunn

## Abstract

Recent developments in understanding how the functional phenotype of the innate immune system is programmed has led to paradigm-shifting views on immunomodulation. These advances have overturned two long-held dogmas: only adaptive immunity confers immunological memory and innate immunity lacks specificity. This work describes the novel observation that innate immune effector cells can be recruited to specific tissues of the body where pathology is present by using a microbial-based immune stimulus that consists of an inactivated pathogen that typically resides or causes infection in that target tissue site. We demonstrate this principle using experimental models of cancer and infection for which different subcutaneously delivered microbial-based treatments were shown to induce the recruitment of immune effector cells to specific diseased organs. Amelioration of disease in a given organ niche was dependent on matching the correct microbial stimulus for the affected organ site but was independent of the nature of the pathology. This observation intriguingly suggests that the immune system, upon pathogen recognition, tends to direct its resources to the compartment in which the pathogen has previously been encountered and would be the most likely source of infection. Importantly, this phenomenon provides a novel means to therapeutically target innate immune effector cells to sites of specific disease localization to potentially treat a wide spectrum of pathologies, including cancer, infection, and chronic inflammatory disorders.

**AUTHOR SUMMARY:** Vaccines that target adaptive immune memory have revolutionized medicine. This study describes a novel strategy that works as a modified innate immune “vaccine” that exploits the trained response of innate immune effector cells to clear pathology in a specific tissue site. Unlike memory of the adaptive immune system, which functions like a lock and key, innate immune memory is more akin to a reflex response – like experienced muscle or neural cells that are changed by a stimulus to respond more efficiently upon re-exposure. This change in behavior through experience is the definition of learning. Our study suggests that this innate immune learning occurs at different levels. Emergency hematopoiesis trains new innate immune cells in the bone marrow to respond quickly and effectively to a non-specific threat; whereas, pathogen-specific training occurs at sites where cells making up the immunologic niche have had interactions with a particular pathogen and have been trained to respond more robustly to it upon re-presentation in the context of a danger signal. The speed with which new immune cells are trained in the bone marrow in response to an imminent microbial threat and their subsequent recruitment to the target organ site where that microbe typically resides suggests there are ways the immune system communicates to coordinate this rapid response that are yet to be fully delineated. These findings provide a novel highly proficient way to harness the potent effector functions of the innate immune system to address a wide range of immune-based diseases.

## INTRODUCTION

In 1899, cancer researcher D’Arcy Power noted: “Where malaria is common, cancer is rare.”[1] For centuries, physicians have observed that acute infection can be associated with spontaneous cancer regression, and it is now well appreciated that the potent immune response to the threat of acute infection can overcome malignancy [2,3]. One of the best known early clinical applications of this observation was developed in the 1890’s by Dr. William Coley who used an inactivated bacterial cocktail, which came to be known as Coley’s Toxin, to treat various types of cancers and achieved success rates similar to modern treatments in many cases [2]. At least 15 epidemiological studies have examined the possible link between infectious disease and cancer, with all but one supporting an association between infection and reduced incidence of neoplasms [2]. Yet, intravesical administration of Bacillus Galmette-Guerin (BCG) for the treatment of high-risk non-muscle invasive bladder cancer is the only microbe-based therapy that is currently part of the approved standard of care [4]. The failure to realize the full potential of this immunotherapeutic approach can be attributed to a number of factors, including lack of sufficient characterization of the immunological mechanism(s) driving the therapeutic outcome and difficulties with obtaining consistent results. However, recent pivotal developments in understanding the basis of what is now coined trained innate immunity, which encompasses changes in the programming of the innate immune response through experience, have far-reaching potential to improve our understanding of the anti-cancer immunological response induced by acute infection [5,6]. This new deeper appreciation of innate immune system adaptation and reprogramming is projected to meaningfully advance the use of microbial-based treatments for not only cancer, but also for many of the increasingly prevalent immune-based diseases we now contend with, including inflammatory bowel disease, allergies and autoimmune disorders [5–7].

We recently characterized the key immune cellular and molecular pathways by which mimicking an acute infection using a microbe-based stimulus markedly reduces the tumor burden in two different lung cancer models [8]. The treatment used was formulated from an inactivated lung pathogen, a derivative of a clinical isolate of *Klebsiella*, which is subcutaneously administered every second day [8]. Efficacy in reducing lung tumor burden was shown to be 1) dependent on the animals having had previous exposure to *Klebsiella* and 2) independent of adaptive immunity [8]. In the work presented here, we describe a newly discovered phenomenon of organ-specific trained innate immune cell recruitment and disease amelioration. We show organ specificity is determined by the organ niche of the bacterial species from which the microbial stimulant is derived. These findings suggest that there are levels of greater complexity of innate immune training that are yet to be fully elucidated. By applying this immunological targeting therapeutically, these findings provide a means to direct trained innate immune effector cells to specific tissue sites of pathology.

## RESULTS

### Microbial-based immunotherapy agents demonstrate anti-cancer efficacy in an organ-specific manner

A subcutaneously injected microbial-based immunotherapy made from an inactivated strain of *Klebsiella*, called QBKPN, was previously tested in a Lewis Lung Carcinoma model [8]. It was observed that mice without prior lung exposure to *Klebsiella* did not experience a reduction in tumor burden with QBKPN treatment, whereas mice with prior exposure to *Klebsiella* did [8]. Here we show that when we administer a similarly formulated immunotherapy based on *Escherichia coli*, called QBECO, there is little relative efficacy in reducing lung cancer burden in the Lewis Lung Carcinoma model, despite all animals likely having exposure to *E. coli* (Fig. 1A). We replicated the previous findings in a different lung cancer model using B16F10 melanoma cells (Supplemental Fig. 1A), which suggests QBKPN’s anti-cancer efficacy is not cancer cell type specific, rather it appears to be applicable to different cancer cell types growing in the lungs. We hypothesized that QBECO may have a therapeutic anti-tumor effect in a compartment likely to be exposed to *E. coli*. We first used an intraperitoneal cancer model since *E. coli* infection is the most common cause of Gram-negative peritonitis [9]. MC-38 adenocarcinoma cells were administered by intraperitoneal injection to establish the cancer [10]. In this model every second day subcutaneous administration of QBECO significantly increased survival compared to vehicle treatment; whereas, QBKPN treatment was less efficacious (Fig. 1B). QBECO treatment also improved survival when pancreatic cancer cells were seeded intraperitoneally (Supplemental Fig. 1B). Thus, QBECO treatment improved survival in animals with cancer in the peritoneal region independent of cancer cell type, corroborating the analogous observation with respect to QBKPN’s anti-cancer efficacy in the lungs. Collectively, these findings suggest there are organ-specific immune effects of these microbial treatments and that these subcutaneously administered products are sampled by immune cells in the targeted organ niche. To exclude the possibility that the efficacy of these microbial-based treatments was a result solely of reduced seeding of injected cancer cells, we tested both QBKPN and QBECO in a chemically-induced colon cancer model in which cancer develops with exposure to DSS/AOM over a 70 day period [11,12]. Because both *Klebsiella* species and *E. coli* are enteric bacteria in mice, both QBKPN and QBECO treatment succeeded in reducing tumor burden in the colon compared to vehicle-treated mice in this model (Fig. 1C). A biodistribution study using *in vivo* imaging of fluorescently labelled QBKPN (Cy5.5-QBKPN) confirmed that the product is systemically distributed (Supplemental Fig. 2A); peak levels in blood were detected at 2 hours after administration and dropped subsequently (Supplemental Fig. 2B).

**Figure 1:**
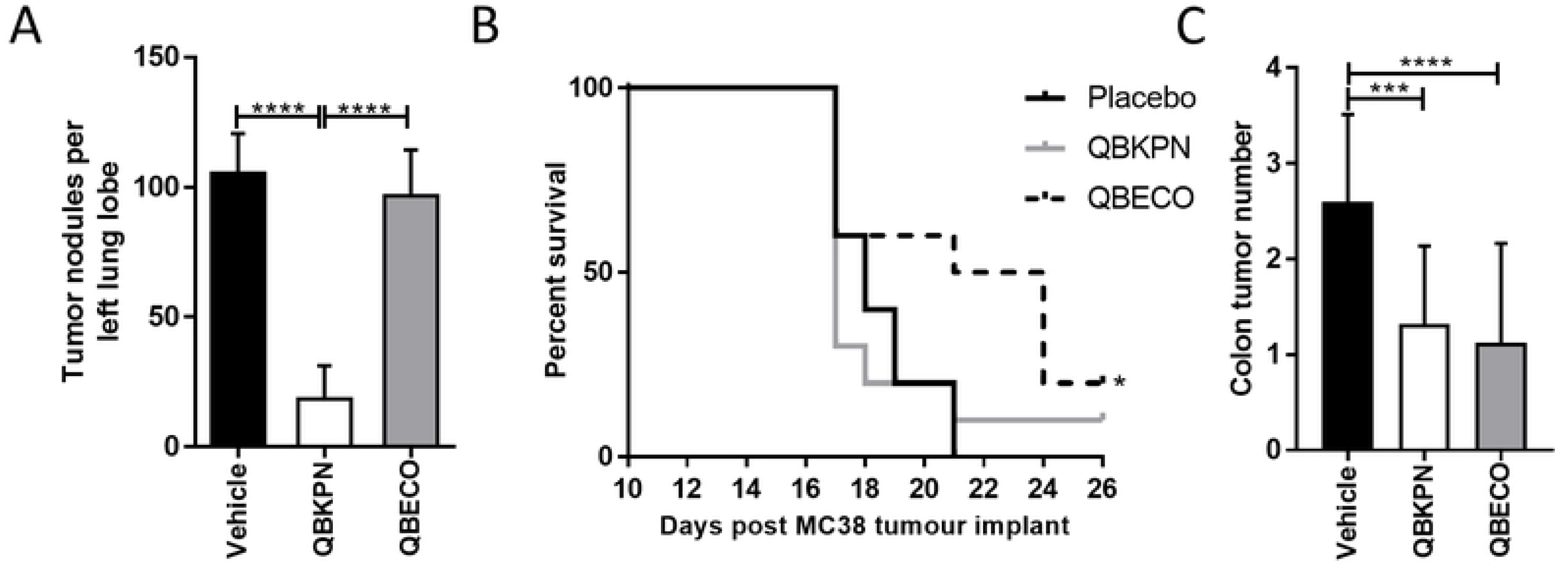
Microbial-based therapies, QBKPN and QBECO, have different organ-specific anti-cancer efficacies. (A) Left lung tumor nodule counts 15 days after mice were seeded with Lewis Lung Carcinoma (LLC)-red fluorescent protein (RFP) cells by tail vein injection and treated with vehicle, QBKPN, or QBECO. n=5 mice per group. (B) Survival curves to day 26 in mice seeded with MC-38 adenocarcinoma cells by intraperitoneal injection and treated with vehicle, QBKPN, or QBECO. n=10 mice per group. (C) AOM/DSS model of spontaneous colon cancer development in mice treated with vehicle, QBECO or QBKPN at day 70. n=19-20 mice per group. For each experiment, mice were given the designated treatment by subcutaneous injection every second day starting 10 days before tumor seeding and continuing throughout the experiment. Data is presented as the mean ± SD. * p < 0.05; ** p < 0.01; **** p < 0.0001, one-way ANOVA with Holm-Sidak’s and Log-rank test.

**Figure 2:**
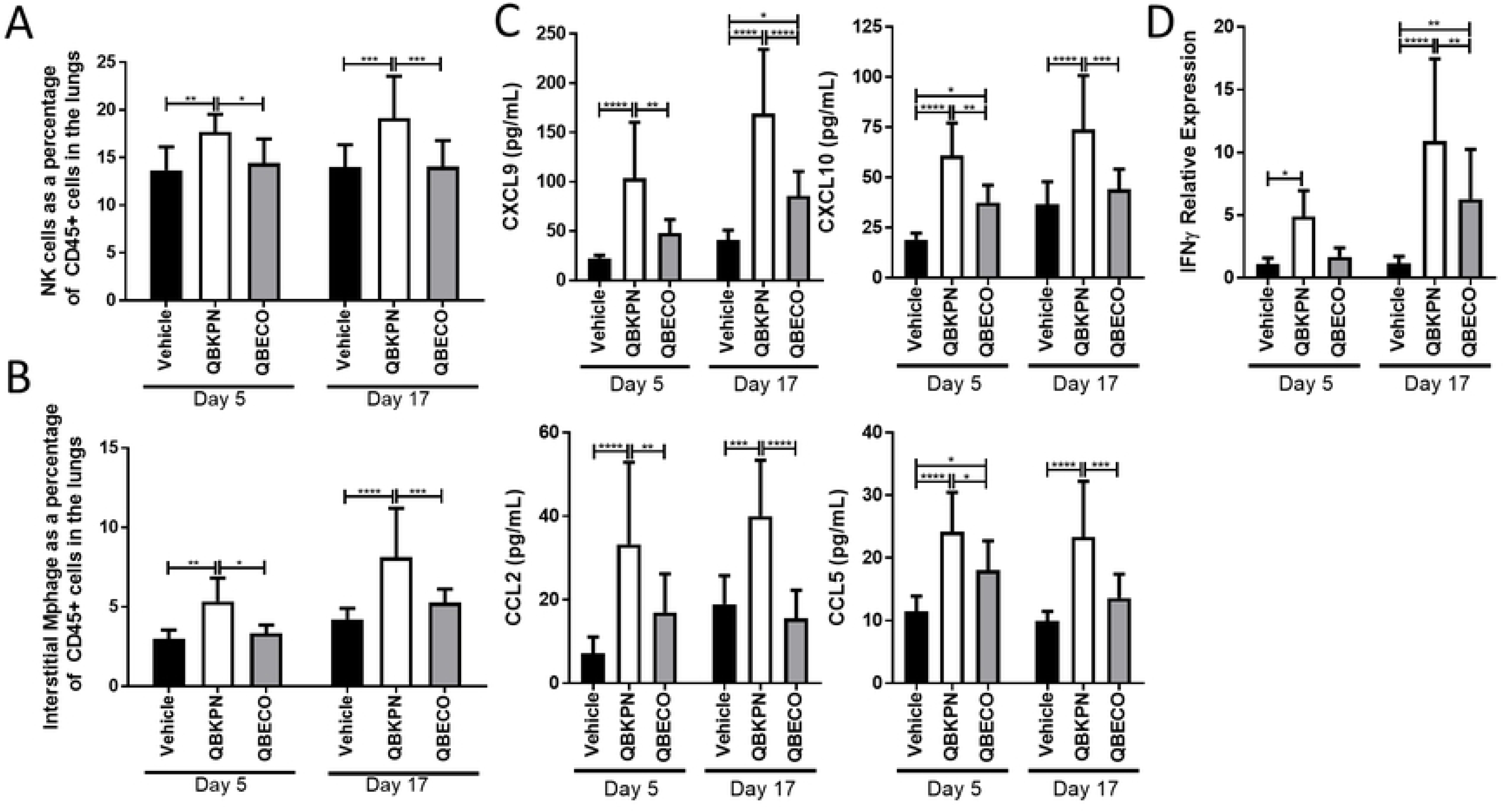
QBKPN, but not QBECO, induces lung specific immune changes in a lung cancer model. **(A,B)** Lung immunophenotyping of mice in the B16F10 lung cancer model at 5 and 17 days post tumor inoculation for **(A)** NK cells and **(B)** interstitial macrophages as a percentage of CD45^+^ cells. Data represents mean ± SD; n = 9-10 mice per group. **(C)** Protein levels in lung homogenates of CXCL9, CXCL10, CCL2 and CCL5 in the B16F10 lung cancer model at 5 and 17 days post tumor inoculation. Cytokines were measured by 31-cytoplex; additional measured cytokines are in Supplemental Fig. 3. Data represents mean ± SD. n=8-10 mice per group. **(D)** IFNγ expression in the lungs tissue in the B16F10 lung cancer model at 5 and 17 days post tumor inoculation as measured by qRT-PCR. Data represents mean ± SD; n=9 to 10 mice per group. In each experiment, mice were treated with vehicle, QBKPN, or QBECO by subcutaneous injections every second day starting 10 days before tumor seeding and continuing throughout the experiment. * p < 0.05; ** p < 0.01; *** p < 0.001; **** p < 0.0001, two-way ANOVA, Tukey’s multiple comparison test.

### Characterizing the lung-specific immune response induced by QBKPN compared to QBECO in mice with lung cancer

Immuno-phenotyping was used to characterize the lung specific immune changes induced by QBKPN compared to QBECO treatment using the murine B16F10 lung cancer model. QBKPN administration increased the percentage of NK cells and interstitial macrophages in the lungs when assessed at both days 5 and 17 after tumor inoculation (corresponding to 15 and 27 days of QBKPN treatment, respectively). In contrast, QBECO treatment did not change the proportion of either NK cells or interstitial macrophages in the lungs relative to vehicle treated control mice (Fig. 2A-B).

To account for the greater recruitment of NK cells and macrophages to the lungs by QBKPN vs. QBECO treatment, lung chemokine concentrations were quantified. Relative to both vehicle control and QBECO, treatment with QBKPN led to higher levels of chemokines important for the recruitment and activation of NK cells and macrophages, including CCL2, CCL5, and IFNγ-inducible chemokines, CXCL9 and CXCL10, at both 5 and 17 days following tumor inoculation (Fig. 2C). IFNγ gene expression was increased with QBKPN treatment compared to QBECO at both timepoints measured (Fig. 2D); however, protein levels of IFNγ were measurably greater at only the day 5 timepoint (Supplemental Fig. 3). QBKPN’s anti-lung cancer efficacy was previously found to be dependent on the activation of the NKG2D pathway, which is fundamental for the detection and elimination of transformed, damaged and infected cells by cytotoxic lymphocytes [8]. QBKPN administration, but not QBECO administration, significantly potentiated the expression of NKG2D ligand, Rae1, in the lungs (Fig. 3A). In parallel, QBKPN treatment led to a more potent induction of molecules that mediate anti-tumor cytotoxicity including, Granzyme A (GzmA), Granzyme B (GzmB) and Perforin 1 (Pfr1), in the lungs of mice with cancer (Fig. 3B-E).

**Figure 3:**
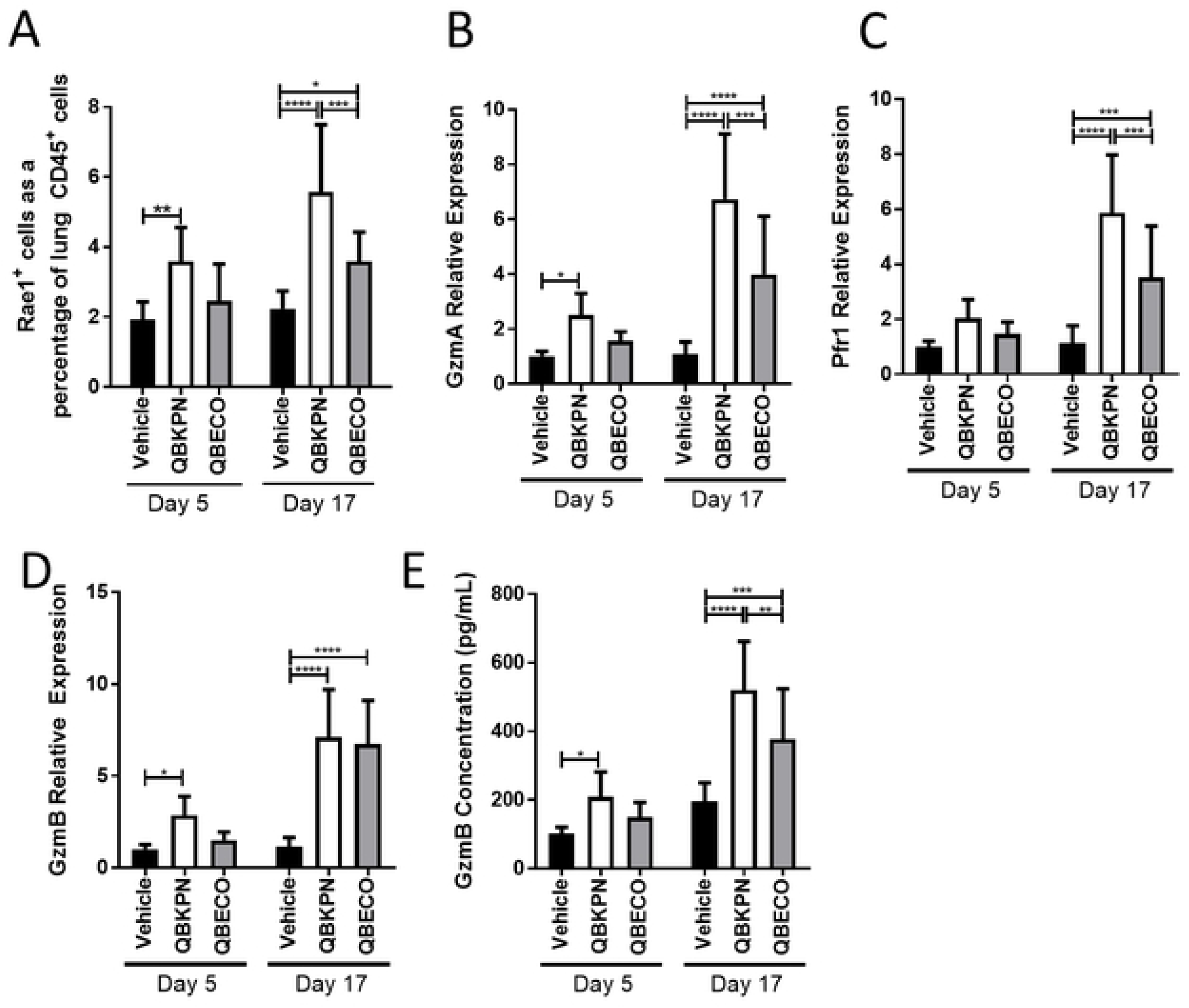
QBKPN, but not QBECO, activates anti-cancer mechanism in lungs. **(A)** Rae1^+^ expression on CD45^+^ cells in the lungs of mice in the B16F10 lung cancer model at 5 and 17 days post tumor inoculation, as assessed by flow cytometry. **(B,C,D)** Gene expression in the lungs of B16F10 cancer bearing mice at 5 and 17 days for **(B)** Granzyme A, **(C)** Perforin 1, and **(D)** Granzyme B. **(E)** Protein level of Granzyme B as measured by ELISA in the lungs of mice in the B16F10 lung cancer model at 5 and 17 days post tumor inoculation. For all studies, data represents mean ± SD. n = 9-10 mice per group. In each experiment, mice were treated with vehicle, QBKPN, or QBECO by subcutaneous injections every second day starting 10 days before tumor seeding and continuing throughout the experiment. * p < 0.05; ** p < 0.01; *** p < 0.001; **** p < 0.0001, two-way ANOVA, Tukey’s multiple comparison test.

### The lung-specific immune response elicited by QBKPN is enhanced by the presence of pathology

The contribution of the presence of a lung pathology to the type and extent of the immune response triggered by QBKPN vs. QBECO was investigated using cancer-free mice. Among the most notable differences in the immune response in healthy animals was the proportion of NK cells recruited to the lungs with QBKPN treatment, which was markedly attenuated in healthy mice compared to lung cancer bearing mice (Fig. 4A vs. Fig. 2A). In contrast, the proportion of interstitial macrophages increased in both healthy mice and mice with cancer (Fig. 4B, Fig. 2B). Another difference in healthy vs. lung cancer bearing mice was the degree to which certain immune responses were sustained. In particular, the expression of IFNγ and Rae1, which decreased or remained the same over time in healthy mice (Fig. 4), increased over time in cancer-bearing mice (Fig. 3). However, similar to cancer-bearing mice, there was greater lung immune stimulation with QBKPN than with QBECO treatment, but this difference was generally less pronounced in healthy mice. The exception to this observation was in the gene expression, but not protein levels, of granzyme B, granzyme A, and perforin, which showed equal increases with 27 days of either QBKPN or QBECO treatment in disease-free animals (Fig. 4G). Collectively, these findings suggest that the presence of pathology dictates the extent, duration and type of lung-specific immune response elicited by QBKPN vs. QBECO treatment.

**Figure 4:**
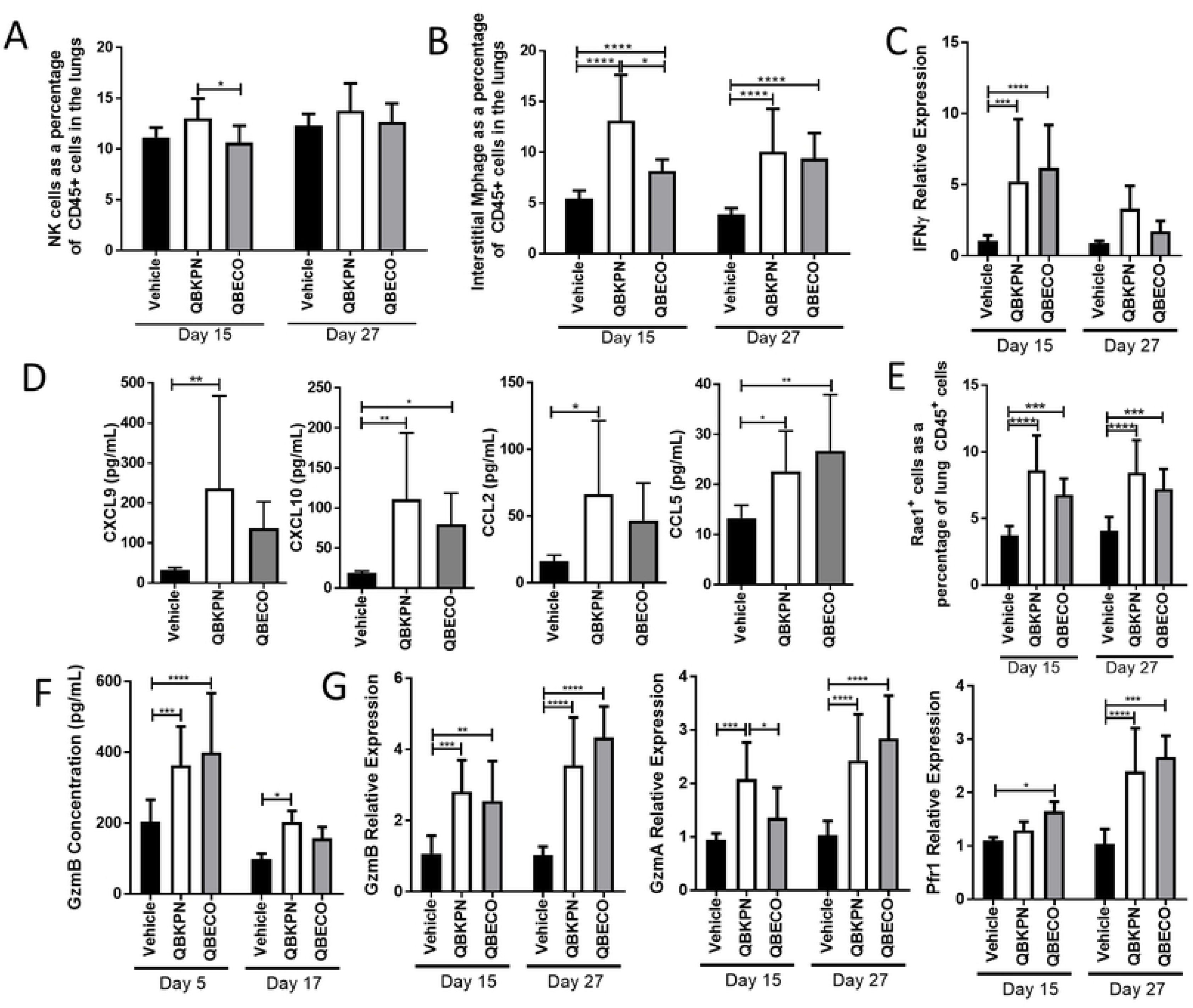
The lung enhanced response to QBKPN is limited in cancer free mice. **(A,B)** Lung immunophenotyping of cancer free mice after 15 and 27 days (which corresponds to the same treatment duration as day 5 and day 17 in the cancer models) of vehicle, QBKPN and QBECO treatment for **(A)** NK cells and **(B)** interstitial macrophages as a percentage of CD45^+^ cells. **(C)** IFNγ expression in the lungs tissue of cancer free mice. **(D)** Cytokine level in lung homogenates for CXCL9, CXCL10, CCL2 and CCL5 in cancer free mice after 27 days of Vehicle, QBKPN and QBECO treatment. Cytokines were measured by 31-cytoplex; additional measured cytokines are in Supplemental Fig. 4. **E)** Rae1^+^ expression on CD45^+^ cells in the lungs of cancer free mice, as assessed by flow cytometry. **F)** Protein levels of Granzyme B as measured by ELISA in the lungs of cancer free mice. **(G)** Gene expression in the lungs of cancer free mice for Granzyme A, Perforin 1, and Granzyme B measured by qRT-PCR. For all studies, data represents mean ± SD. n = 9-10 mice per group. * p < 0.05; ** p < 0.01; *** p < 0.001; **** p < 0.0001, two-way ANOVA, Tukey’s multiple comparison test **(A,-C, E-G)**, one-way ANOVA with Holm-Sidak’s post-hoc test **(D)**.

### Both microbial-based treatments, QBKPN and QBECO, stimulate an initial acute immune response that results in emergency hematopoiesis

The nature of the initial systemic immune response elicited by QBKPN and QBECO following a single subcutaneous injection was first assessed in healthy animals to exclude the impact of organ pathology on their different organ-specific effects. The kinetics and magnitude of the systemic acute-immune response were similar after administration of the two microbial-based products (Fig. 5A-C). Both treatments induced a rapid induction of systemic inflammatory cytokines that were detectable in the serum within 5 hours and returned to baseline levels by 12 to 24 hours after administration (Fig. 5A; Supplemental Fig.5). Similarly, both QBKPN and QBECO increased the number of neutrophils and Ly6C^HI^ monocytes in circulation within 5 hours. (Fig. 5B-C).

**Figure 5:**
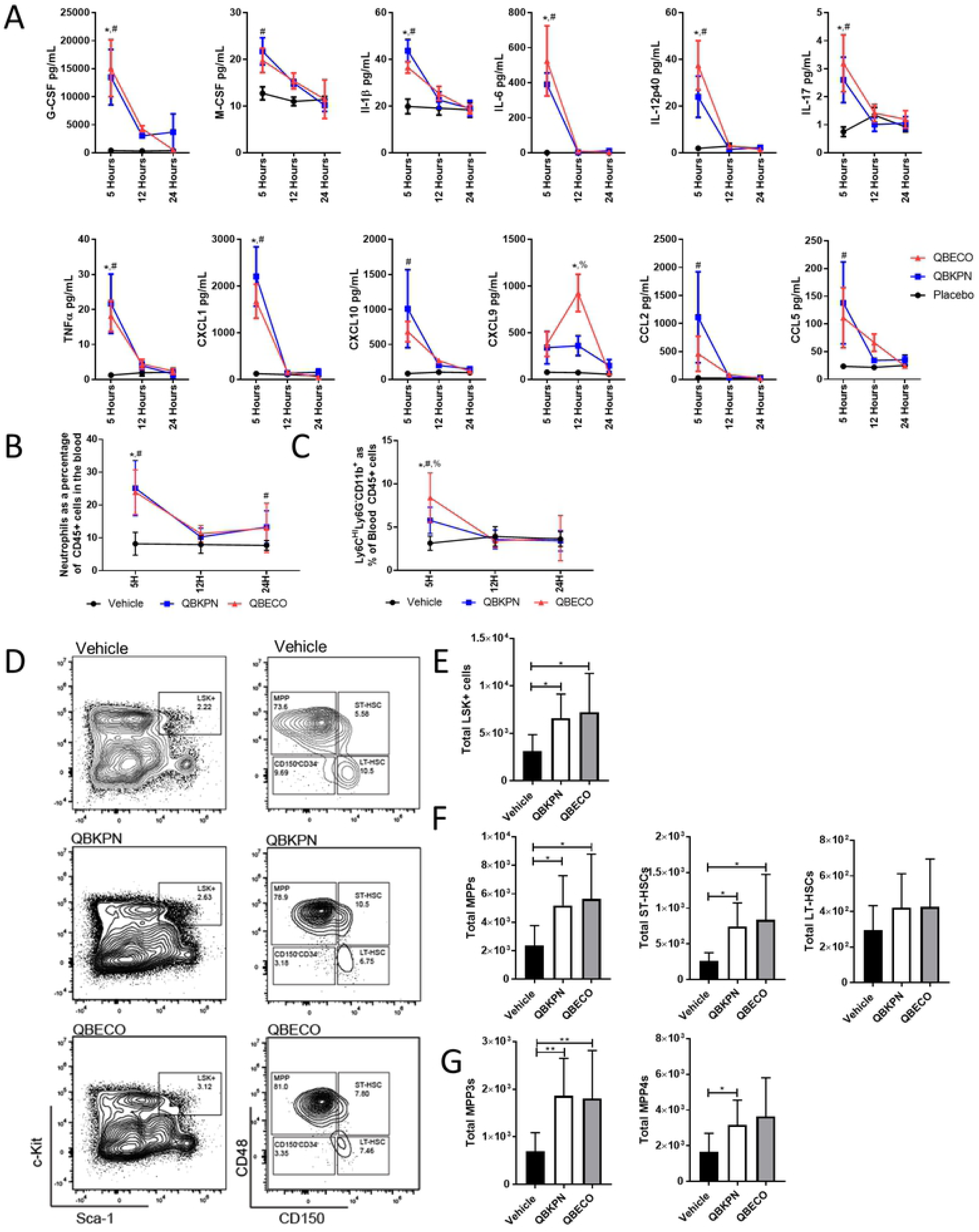
QBKPN and QBECO create an early cytokine response and stimulate emergency hematopoiesis. **(A)** Cytokine measurements in the serum 5, 12 and 24 hours after treatment with vehicle, QBKPN or QBECO by subcutaneous injection. Cytokines were measured by 31-cytoplex; additional measured cytokines are in Supplemental Fig. 5. N=9 to 10 mice per group; mean ± SEM. Data from 5 hours in vehicle and QBKPN treated mice has been previously published[8]. **(B)** Neutrophils (Ly6G^+^Ly6C^-^) and **(C)** Monocytes (Ly6C^HI^Ly6G^-^CD11b^+^) as a percentage of CD45^+^ blood cells. n=10 mice per group. Mean ± SD shown. **(D)** Representative flow cytometry plots of the bone marrow 24 hours after mice were treated with a single dose of subcutaneous vehicle, QBKPN and QBECO for LSK+ (Lin^-^Sca-1^+^c-Kit^+^) and for multipotent progenitors (MPPs; CD150^-^CD48^+^), short-term HSCs (ST-HSC; CD150^+^CD48^+^), and long-term HSCs (LT-HSC; CD150^+^CD48^-^)cells. Cell counts of **(E)** LSK^+^ cells, **(F)** MPPs, ST-HSC, and HT-HSC, and **(G)** myeloid lineage MPPs (MPP3s; MPP^+^CD34^+^Flt3^-^) and lymphoid lineage MPPS (MPP4s; MPP^+^CD34^+^Flt3^+^) at 24 hours after a single subcutaneous dose of vehicle, QBKPN and QBECO. n=10 mice per group. Mean ± SD shown. **(A-C)**\* p < 0.05 for QBKPN vs Vehicle; ^#^ p < 0.05 for QBECO vs Vehicle; ^%^ p < 0.05 for QBKPN vs QBECO, two-way ANOVA, Tukey’s multiple comparison test. **(E-G)** * p < 0.05; ** p < 0.01; *** p < 0.001; **** p < 0.0001, one-way ANOVA with Holm-Sidak’s post-hoc test.

Changes in the hematopoietic stem cell (HSC) populations were investigated to assess how a single injection of either QBKPN or QBECO affected hematopoiesis and the mobilization of new immune cell recruitment. Twenty-four hours after treatment, both QBKPN and QBECO similarly expanded the HSC progenitor population in the bone marrow, characterized as lineage negative c-Kit^+^Sca^+^ (LSK^+^) (Fig. 5D-E). In particular, both microbial products increased the production of short-term HSC (ST-HSC; characterized as LSK^+^CD150^+^CD48^+^) and multipotent progenitors (MPPs; LSK^+^CD150^-^CD48^+^); Fig. 5G. Both myeloid (MPP3; LSK^+^CD150^-^CD48^+^CD34^+^Flt3^-^) and lymphoid (MPP4; LSK^+^CD150^-^CD48^+^CD34^+^Flt3^+^) MPP lineages were similarly increased at this time point with QBKPN and QBECO treatment.

### In lung cancer bearing mice, QBKPN-mediated myelopoiesis induces recruitment of trained innate immune cells to clear pathology

Microbial-based stimulation of myelopoiesis is considered to be an important component of trained innate immunity required to deal with the threat of infection [13]. To determine whether this process is also involved in mediating QBKPN’s ability to clear cancer in the lungs, the modulation of myelopoiesis by QBKPN and QBECO was investigated in the context of cancer in the lungs. Mice repeatedly treated with QBKPN, but not mice treated with QBECO, had sustained increases in the number of HSC (LSK^+^ cells) compared to vehicle-treated control mice when the bone marrow niche was assessed 18 days after B16F10 cancer cell administration seeded to the lungs (Fig. 6A). This QBKPN-mediated increase in progenitors favoured MPPs and ST-HSCs (Fig. 6B). Within the MPP population, repeated treatment with QBKPN in lung cancer-bearing mice resulted in a shift towards MPP3 generation (Fig. 6C). In contrast, QBECO treatment had little effect in modulating the HSC populations in lung-cancer bearing mice when assessed at this later time point (Fig. 6A-C). Both QBKPN and QBECO treatment induced large increases in bone marrow neutrophil numbers; however, only QBKPN was able to sustain an increased number of bone marrow monocyte numbers (Fig. 6D). To test whether bone marrow monocytes in mice treated with these microbial-based therapies were functionally different, we isolated bone marrow monocytes from B16 lung tumor bearing mice treated with vehicle, QBKPN or QBECO and stimulated them with LPS *in vitro*. Bone marrow monocytes from mice treated with QBKPN or QBECO released significantly more IL-1β relative to those treated with vehicle. (Fig. 6E).

**Figure 6:**
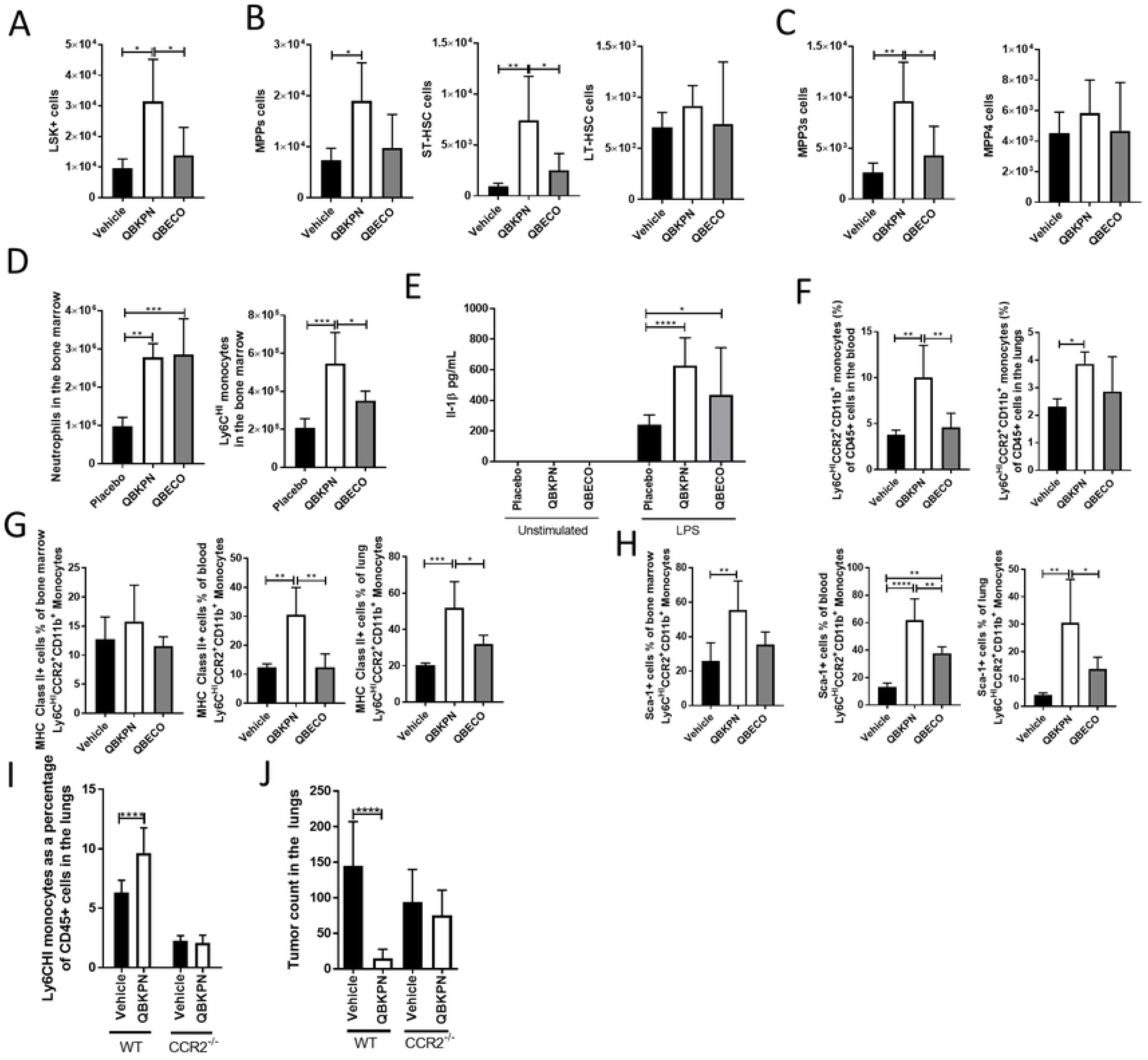
Repeated administration in a cancer environment promotes myelopoiesis and induction of trained monocytes. **(A-C)** Cell counts of LSK^+^ cells, MPPs, ST-HSC, HT-HSC, CD150^+^CD45^-^LSK^+^ cells, MPP3s (MPP^+^CD34^+^Flt3^-^) and MPP4s (MPP^+^CD34^+^Flt3^+^) in B16F10 bearing mice 18 days after tumor inoculation, and administered vehicle, QBKPN and QBECO. n=5 mice per group. **(D)** Total number of neutrophils and Ly6C^HI^ monocytes in the bone marrow in the day 18 B16F10 lung cancer model. n=5 mice per group. **(E)** IL-1β levels in the supernatant of bone marrow monocytes originally isolated from mice in the day 17 B16F10 lung cancer model and subsequently stimulated for 24 hours with LPS (1ng/mL) or left unstimulated. n = 5 mice per group. No IL-1β production was detected in the unstimulated samples. **(F)** Ly6C^HI^CCR2^+^CD11b^+^ monocytes as a percentage of all CD45^+^ cells in the bone marrow, blood, and lungs at day 18 in the B16F10 cancer model. n=5 mice per group. **(G)** MHC Class II expression on Ly6C^HI^CCR2^+^CD11b^+^ monocytes in the blood, and lungs at day 18 in the B16F10 cancer model. n=5 mice per group. **(H)** Sca-1 expression on Ly6C^HI^CCR2^+^CD11b^+^ monocytes in the bone marrow, blood, and lungs at day 18 in the B16F10 cancer model. n=5 mice per group. **(I)** Monocytes (Ly6C^HI^CD11b^+^Ly6G^-^) and **(J)** tumor burden, in the lungs at day 15 in B16F10 cancer bearing wildtype (WT) and CCR2 knockout (CCR2^-/-^) mice administered with QBKPN and vehicle. n = 8 mice per group. In each experiment, mice were treated with vehicle, QBKPN, or QBECO by subcutaneous injections every second day starting 10 days before tumor seeding and continuing throughout the experiment. Mean ± SD shown. * p < 0.05; ** p < 0.01; *** p < 0.001; **** p < 0.0001, one-way ANOVA with Holm-Sidak’s post-hoc test **(A-H)**, Student’s *t-*test **(I-J)**.

When we investigated systemic immune cell populations in the lung cancer model at Day 18, we found QBKPN, but not QBECO, treatment resulted in an increase in Ly6C^HI^CCR2^+^CD11b^+^ monocytes in circulation and lungs (Fig. 6F). These monocytes were characterized as being positive for MHC class II and Sca-1 (Fig. 6G-H), markers associated with transcriptional programming in the bone marrow by microbial stimulation [14]. Ly6C^HI^ monocytes are recruited to sites of infections through CCR2/CCL2 signalling [15,16]. The important contribution to anti-lung cancer efficacy of QBKPN-mediated recruitment of these monocytes was demonstrated in mice genetically deficient in CCR2, in which QBKPN treatment failed to recruit Ly6C^HI^ monocytes to the lungs and lost its anti-cancer efficacy (Fig. 6I-J).

### Demonstration of the broader application of immune modulation by organ-specific microbial-based therapies to treat immune-related diseases

The ability to target trained innate immune cells to specific organ sites of pathology by the strategic selection of the correctly matched microbial-based therapy has potential to treat a broad range of immune-related diseases. Adding to the evidence provided for this novel approach for cancer using specific gram-negative pathogens targeting the lungs, peritoneal cavity and GI tract, we next tested the ability to target the skin using a microbial-based treatment (QBSAU) made from a clinical isolate of *Staphylococcus aureus*, a Gram-positive organism that is a common source of skin infection. Treatment with QBSAU, but not QBKPN or QBECO, was effective in reducing B16F10 melanoma subcutaneous tumor growth (Supplemental Fig. 6A). In contrast, QBSAU had little relative efficacy in improving outcomes for animals with cancer in the lungs or peritoneal compartment (Supplemental Fig. 6B-C).

To demonstrate the broader application of this immunotherapeutic approach, we tested the three different microbial-based treatments, QBKPN, QBECO, and QBSAU, in three infection models affecting the lungs, peritoneal cavity and skin. Pathogens different from those used to make the microbial-based treatments were used for establishing organ-specific infections to avoid the possibility that the treatments were functioning like classical vaccines through the induction of an adaptive immune response to a specific pathogen. In the *Streptococcus pneumoniae* lung infection model, out of the three microbial-based treatments used, QBKPN performed the best in reducing the bacterial burden in the lungs (Fig. 7A). In corollary, the intraperitoneal *Salmonella enterica* infection model showed that QBECO treatment had the greatest efficacy in reducing the peritoneal bacterial burden as assessed in the spleen (Fig. 7B). Lastly, in the *Pseudomonas aeruginosa* skin infection model, QBSAU treatment was the most effective in reducing the skin bacterial burden (Fig. 7C). The ability of these microbial-based therapies to ameliorate disease was independent of the species of pathogen causing infection. QBKPN reduced bacterial burden in the context of both *S. pneumoniae* and *P. aeruginosa* (Supplemental Fig. 7) infections.

**Figure 7:**
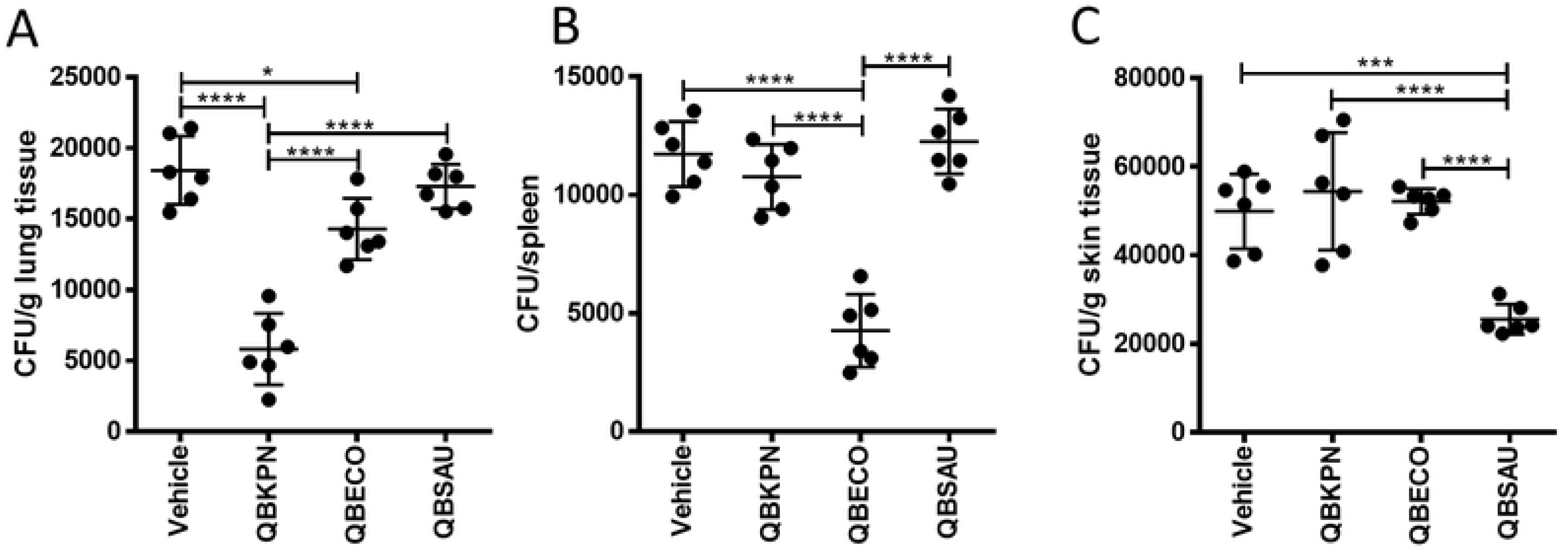
Microbial-based treatment protect against infections in an organ-specific manner. **(A)** Lung bacterial counts days 3 days after mice were challenged with *Streptococcus pneumoniae* by intratracheal instillation and treated with vehicle, QBKPN, QBECO or QBSAU. **(B)** Spleen bacterial counts in mice challenged with *Salmonella enterica (typhimurium)* by intraperitoneal injection and treated with vehicle, QBKPN, QBECO or QBSAU. **(C)** Skin bacterial counts 3 days after mice were challenged with a subcutaneous infection of *Pseudomonas aeruginosa* and treated with vehicle, QBKPN, QBECO or QBSAU. In each experiment, treatments were given by subcutaneous injections every second day as described in the Material and Methods. n = 6 mice per group. Data is presented as mean ± SD. * p<0.05; *** p < 0.001; **** p < 0.0001, one-way ANOVA with Holm-Sidak’s post-hoc test.

## DISCUSSION

Microbial exposure is a double-edged agent of health and disease. Infection was once the leading cause of morbidity and mortality, but with significant improvements in hygiene and the advent of population-based vaccination and antibiotics, the 20^th^ Century saw a dramatic decrease in the incidence of acute infection and infection mortality and a concomitant rise in allergies, chronic immune-related disorders [17,18] and certain cancers [19] in developed countries. This observation has become popularly known as the hygiene hypothesis, which postulates that a lack of appropriate training of the immune system by microbial stimulation, particularly early in life, is linked to the development of immune pathologies. This hypothesis has different variations, some postulating that exposure to disease causing agents is less important than early life exposure to “friendly” or symbiotic microbes [20]. The truth likely encompasses an integration of both concepts. Evidence for the hygiene hypothesis has primarily come from epidemiological studies (reviewed in [17,18,21,22]), but there is direct experimental evidence that immune education by microbial stimulation early in life is important for setting the tone for immune regulation throughout [23]. This has spurred an area of avid research focused on the use of microbes and microbial components as potential new treatment modalities for the many serious immune-related diseases growing in incidence and prevalence [5,24,25]. In this work, we describe what we believe to be a seemingly unknown phenomenon of the existence of immunological memory contained within specific regional niches for microbes that normally inhabit or infect it. We were able to successfully target pathology in the lungs, peritoneal cavity, GI tract and skin via repeated subcutaneous administration of microbial-based immunotherapies made from inactivated bacterial species that are endogenous in each of the respective target organs. The efficacy of a *Klebsiella*-based treatment to clear lung cancer was previously found to be dependent on animals having prior lung exposure to *Klebsiella* and independent of adaptive immunity [8]. Our studies show that the choice of microbe used to formulate the immunotherapy is critically important to appropriately direct innate immune effector cells to the site of pathology. The lack of consistency and high variability in outcomes with microbial-based treatments to date may partly be due to a mismatch between the microbial species from which the treatment is derived and the past microbial exposure or “training” of the patient’s immune system in the targeted organ. Fig. 8 summarizes the sequence of events we have determined leads to the clearance of pathology in a targeted organ by systemically administering a microbial-based stimulus derived from a pathogen that is a common cause of infection in that organ niche.

**Figure 8:**
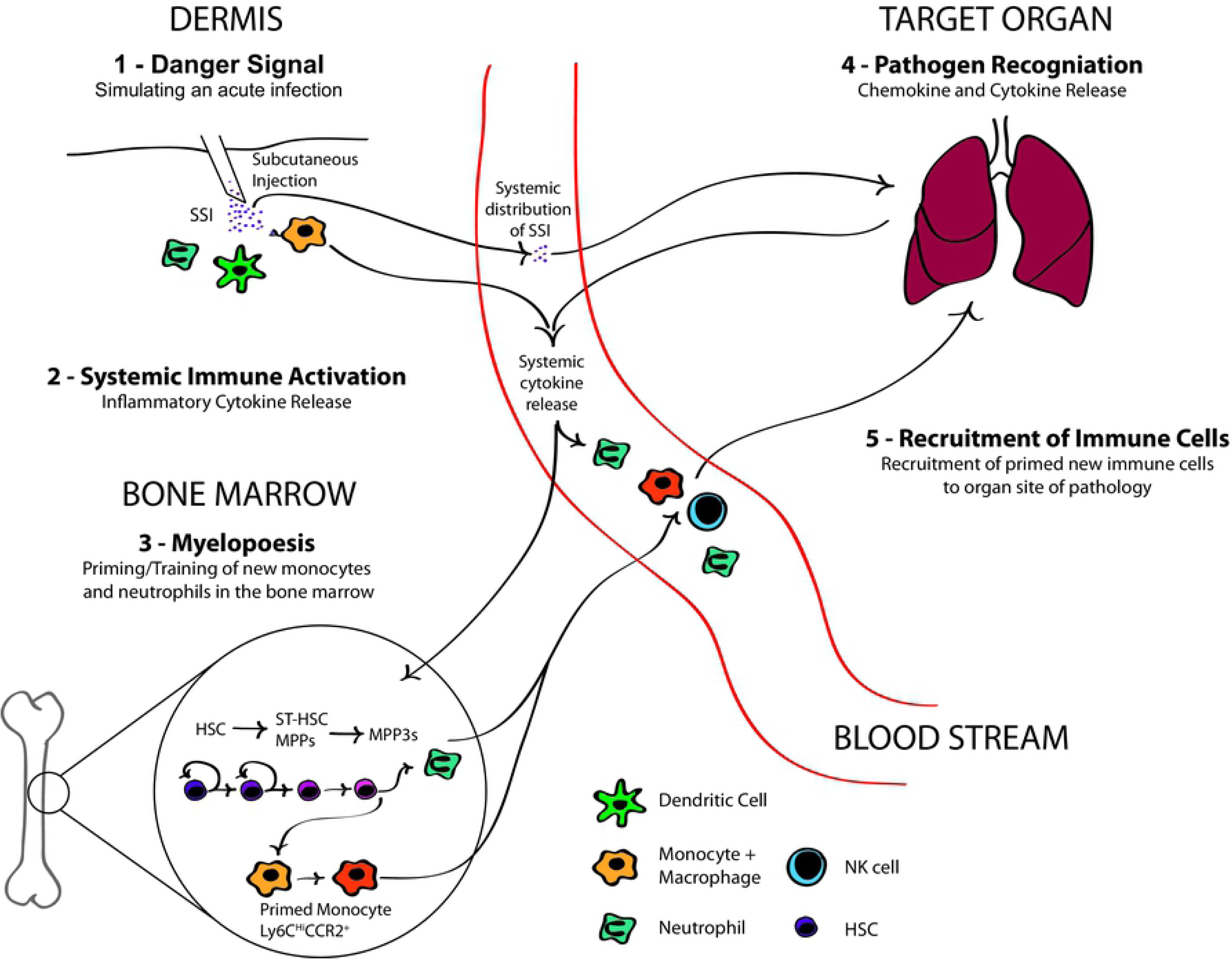
Summary of the mechanism leading to organ-specific immune activation by systemically delivered microbial stimulus. A subcutaneously delivered microbial stimulus elicits a systemic immune response that leads to myelopoeisis and training of new immune cells in the bone marrow to deal with an acute threat. Recognition of the microbial components in the context of an immunological threat in the target organ in which it is a part of the microbiome as well as a common source of infection induces the local production of chemokines that recruit the newly trained immune cells. The end result is an efficient immune response to the pathology present in that tissue.

Two long-held dogmas have recently been brought into question with paradigm-shifting advances in our understanding of innate immunity: 1) immunological memory is the domain of the adaptive arm of the immune system, and 2) innate immunity lacks specific memory [5,6,26,27]. The treatment strategy we employed is rooted in both the trainability of the innate immune system and its more nuanced organ specificity, which is intrinsically different from that of the adaptive immune system [26]. We demonstrated that subcutaneous administration of our microbial-based treatments resulted in myelopoiesis-associated training of monocytes in the bone marrow within 24 hours. Intraperitoneal administration of a prototypical agonist for innate immune training, β-glucan, was first shown to act on HSC to expand myeloid cell progenitors, and this effect on HSC improved the physiological response to subsequent challenge with LPS and also protected mice from chemotherapy-induced myelosuppression [13]. This work by Mitroulis and colleagues established that microbial modulation of myelopoiesis is a fundamental component of trained immunity [13]. Kaufmann and colleagues had come to a similar conclusion in parallel using BCG in a series of elegant experiments that demonstrated BCG administration induced myelopoiesis at the expense of lymphopoiesis and the production of epigenetically modified macrophages that were better able to deal with virulent M. tuberculosis [28]. However, their intent was to demonstrate the longer-term benefits of innate immune conditioning, enabling it to deal more effectively with challenges it will face in the future. We have shown that this innate immune training in the bone marrow can be therapeutically harnessed to deal with a broad spectrum of imminent or present immunological threats by targeting the recruitment of these cells to a specific organ.

Askenase and colleagues discovered that the priming of monocytes in the bone marrow in response to infection was mediated by the production of IFNγ by NK cells, and this proactive developmental education promoted their effector functions in the infected tissues [14]. Microbial-induced innate immune conditioning in the bone marrow is governed by secreted inflammatory signals, including IL-1β, GM-CSF, IL-12 and IFNγ, and largely lacks specificity [13,14]. As we demonstrated, primed monocytes, once recruited to the tissues, can potently respond to a range of dangers. This is consistent with studies demonstrating that BCG vaccination offers non-specific protection against non-mycobacterial infections that is independent of adaptive immune function [29]. In our study, we demonstrated that the signal for targeting immune cells newly released from the bone marrow came from the organ where direct interaction with the microbial species occurs. This organ-specific recognition of the microbial-based treatments was pivotal for the local production of chemokines necessary to its recruitment of trained innate immune effector cells. This is in line with the findings of Olszak *et al.* who demonstrated that early life microbial exposure modulated the recruitment of invariant NKT cells to mucosal sites later in life [23], however, the mechanism by which this microbial regulation is instituted was not defined.

It makes intuitive sense that there exists a means by which the innate immune system directs its resources to the most likely source of infection upon detecting microbial components that are presented as an acute threat. We propose this organ-specific microbial recognition reflects a more sophisticated level of immunological education than the non-specific training that initially occurs in the bone marrow. It is reminiscent of a similar phenomenon observed in invertebrates, which lack adaptive immunity, yet show a stronger immune response to bacterial strains previously encountered compared to strains to which they are naïve [26]. It was previously demonstrated that a *Klebsiella*-based immunotherapy was effective in reducing lung cancer burden in Rag2 deficient mice that are largely devoid of classical adaptive immune function [8]. The contribution of adaptive immunity to the observed microbial mediated organ-specificity cannot be ruled out; however, we believe it is unlikely the primary driver of this organ specific memory. Cells not conventionally considered to be part of the immune system, such as epithelial and endothelial cells that function as the interface between the host and its microbiota, are also capable of forming long-term memories. Acute inflammation was elegantly shown to functionally change epithelial stem cells to respond to such subsequent assaults with improved barrier restoration [30]. On the other hand, barrier dysfunction can also develop through reprogramming of epithelial cells in response to allergic inflammation [31]. This epithelial inflammatory memory, similar to innate immune training [5], is mediated at the level of epigenetic programming [30]. In our study, the presence of pathology in the targeted tissue further potentiated this organ specific immune response to the microbial stimulus, and the nature of the pathology dictated the effector functions executed by the recruited primed innate immune cells.

The therapeutic potential of trained innate immunity has yet to be fully realized. Recent advances in understanding the basis of innate immune programming has re-ignited research into the application of microbial modulation of immune function for the prevention and treatment of disease. The discovery of tissue-specific microbial memory that can be therapeutically exploited to direct trained innate immune effector cells to sites of pathology has the potential to change the way diseases rooted in immune dysfunction are treated. In addition to treating various cancers and infections, proof-of-principle experiments conducted to date show promise for this approach for the treatment of asthma [32], COPD [33], and inflammatory bowel disease [34]. The clinical application of this immunotherapeutic strategy has been promising thus far, based on Phase 1 and 2 studies in non-small cell lung cancer [8], ulcerative colitis [35], and Crohn’s disease [36]. Further research is now needed to fully elucidate the detailed immunological mechanisms of the organ specific innate immune response to microbial threat and its therapeutic application.

## MATERIALS AND METHODS

### Animals

Female C57BL/6J mice (aged 6-10 weeks) were sourced from Jackson Laboratories (Bar Harbor, ME, USA) and Envigo (Livermore, CA, USA). Female CCR2-deficient mice (B6.129S4-Ccr2<tm1lfc>/J) were purchased from Jackson Laboratories. Mice were acclimatized and housed for at least one additional week prior to the experimental studies and were contained in environmentally controlled conditions with a 12:12 hour light/dark cycle for the duration of the study.

### Ethics Statement

All animal care procedures and experimental protocols were approved by the Institutional Animal Care and Use Committee (IACUC) of Dartmouth College (protocol 1141). All animal care procedures and experimental protocols were performed in accordance with applicable international, national, and/or institutional guidelines, including NIH and USDA policies on the care and use of animals in research and teaching. All experimental protocols involving animals performed at Qu Biologics were reviewed and approved by their Animal Care Committee under AUP-201001-001 and were conducted in adherence to the Canadian Council on Animal Care’s (CCAC) policies and guidelines.

### Microbial-based treatments

QBKPN is a proprietary investigational immunotherapeutic formulated from an inactivated clinical lung isolate of *Klebsiella variicola* acquired from a patient with pneumonia [8,32,33]. QBECO is a similarly manufactured proprietary investigational immunotherapeutic made from an inactivated clinical isolate of *Escherichia coli* from a patient with bloody diarrhea. QBSAU is a proprietary investigational immunotherapeutic made from an inactivated clinical isolate of *Staphylococcus aureus* acquired from a patient with an acute subcutaneous abscess. QBKPN, QBECO and QBSAU are suspended in physiological saline, with or without 0.45% phenol as a preservative, and were supplied by Qu Biologics, Vancouver, Canada. The vehicle control used was physiological saline, with or without 0.45% phenol. For experiments performed using only a single dose, 30 μL of the treatment solution was subcutaneously injected into the lower right abdomen. For longer-term experiments involving multiple treatments, subcutaneous injections (into the animal’s skin folds) were administered every second day, rotating among the lower right abdomen, upper right chest, upper left chest, and lower right abdomen of the animal. For cancer model experiments in which the cancer cells were injected into the animal to seed, treatment starting 10 days before tumor cell inoculation (day 0) and every second day throughout the experiment except for the day of tumor inoculation, as previously described [8]. For experiments performed using the DSS/AOM cancer model and naïve mice, the microbial-based treatments were given every second day throughout the experiment. For the infection models, treatment started 14 days before bacterial challenge (day 0) and was given every second day throughout the experiment, except for the day of bacterial challenge.

### Cancer cell injection models

B16-F10 melanoma cells (ATCC CRL-6475, Manassas, VA, USA), red fluorescent protein (RFP)-tagged Lewis lung carcinoma (LLC-RFP) cells (AntiCancer Inc.), MC-38 adenocarcinoma cells (gift from Dr. Jeff Schlom, National Cancer Institute), and Panc02 pancreatic cancer cells (gift from Professor Dieter Kabelitz, Institute of Immunology, Kiel University) were cultured in RPMI medium supplemented with 10% FBS. Cancer cells were harvested and resuspended in PBS before implantation. Cells were confirmed to be free of Mycoplasma infection using the Hek Blue Mycoplasma Detection Kit from Invivogen.

For the lung cancer studies, cells suspended in 100 ul of PBS were injected intravenously by tail vein injection using 2 × 10^5^ B16-F10 cells and 4 × 10^5^ LLC-RFP cells. Tumor burden was assessed at day 15 post-tumor inoculation by enumerating the visible surface metastatic lung lesions; LLC-infiltrated lungs were stained with Bouin’s solution (Sigma-Aldrich, St. Louis, MO, USA) to provide contrast.

For the intraperitoneal cancer survival studies, MC-38 cells or Panc02 cells were injected intraperitoneally at 2 × 10^5^ cells in 100 µL PBS. Mice were monitored daily and euthanized when they reached humane endpoint criteria (i.e., appearance, posture/gait, abdominal distension, behavioural and facial signs of pain).

For the skin cancer study, B16-F10 cells were injected subcutaneously at 1 × 10^5^ cells in 100 µL PBS. Tumor length, width, and height were measured by digital calipers and the tumor burden was calculated using these measurements every second day.

### DSS/AOM model

Mice were injected intraperitoneally on day 0 with azoxymethane (AOM; Sigma, Kawasaki, Kanagawa Prefecture, Japan), at 8 mg/kg body weight, 10 days after treatment started. After AOM injections, mice received 3 cycles of dextran sodium sulphate (DSS) (MP Biomedicals, Santa Ana, CA, USA) treatment spaced 3 weeks apart. In each cycle, DSS was administered in the drinking water *ad libitum* for 5 days at a final concentration of 2.5% weight/volume. Mice were euthanized at the experimental endpoint 70 days after AOM injection.

### Biodistribution Imaging of fluorescently labelled QBKPN

The National Research Council (NRC) of Canada performed the *in vivo* biodistribution studies. Labelling of the microbial-based product, QBKPN, was carried out using Cy5.5 NHSester (GE Healthcare Life Sciences). All optical imaging experiments were performed using a small-animal time-domain eXplore Optix MX2 pre-clinical imager, and images were analyzed or reconstructed as fluorescence concentration maps using ART Optix Optiview analysis software 2.0 (Advanced Research Technologies, Montreal, QC). The pharmokinetic studies were conducted by collecting blood samples, approximately 0.08 mL, from the submandibular vein at selected time points (from 10 minutes to 96 hours after administration) using three mice.

### Flow Cytometry

Anti-mouse antibodies CD16/32, CD45 (30-F11), CD4 (GK1.5), CD8 (53-6.7), NK1.1 (PK136), Ly6G (1A8), Ly6C (HK1.4), F4/80 (BM8), CD11b (M1/70), MHC class II (M5/114.15.2), Rae1 (CX1), CD64 (X54-5/7.1), CD24 (M1/69), CD11c (N418), CCR2 (SA203G11), Sca1 (E13-161.7), CD150 (TC15-12F12.2), CD48 (HM48-1), CD34 (RAM34), Flt3 (A2F10) and c-kit (2B8) were sourced from BioLegend (San Diego, CA, USA) and eBioscience (Waltham, Massachusetts, USA). For HSC determination, lineage cells were stained by Mouse Lineage Antibody Cocktail (Catalog Number 561301, BD Biosceince). Lungs were harvested and dissociated into a single cell suspension using the Milltenyi Biotec (San Diego, CA, USA) mouse lung dissociation kit and gentleMacs tissue dissociator. Bone marrow cells were harvested from both femurs and tibias. Red blood cells were removed from blood, lung and bone marrow samples using Red Blood Cell Lysis Buffer (Biolegend) before staining. Cell viability was assessed using the live/dead fixable violet cell stain kit (ThermoFisher Scientific) and for HSC sample analysis, CD16/32 was blocked using TruStain fcX (BIoLegend). Flow cytometry was performed on a CytoFlex (Bechman Coulter, Indianapolis, IN, USA). Data were analyzed using FlowJo software (version 10.2, Ashland, OR, USA). Lung immune cell profiles were created using a protocol modified from Yu *et al.*[37] while bone marrow hematopoietic stem cells populations were gated as defined by Kaufmann *et al.* [28].

### Immune mediator profiling

Lung tissue samples were homogenized using TissueLyser LT (Qiagen, Hilden, Germany) in 0.5mL of 1% Tween-20 (Sigma-Aldrich, St. Louis, MO, USA) solution. Serum and lung immune factors were analyzed by multiplex technology (performed by Eve Technologies, Calgary, AB, Canada) using a 31 cytokine/chemokine/growth factor kit (Millipore, St Charles, MO, USA). The assay was run on a Bio-Plex™ 200 system (Bio-Rad Laboratories, Inc., Hercules, CA, USA). Data from 5 hours in vehicle and QBKPN treated mice has been previously published [8].

### Gene expression

The right lung post-caval lobe was homogenized by a TissueLyser LT (Qiagen, Toronto, ON, Canada). RNA was isolated using the PureLink RNA Mini Kit (Life Technologies, Carlsbad, CA, USA) and reverse transcribed into cDNA using the iScript cDNA Synthesis Kit (Bio-Rad). Gene expression was quantified by quantitative RT-PCR using a StepOnePlus RT-PCR machine (Applied Biosystems, Foster City, CA, USA), TaqMan Fast Advanced Master Mix (Applied Biosystems) and Taqman probes (Applied Biosystems) for granzyme A (Mm01304452_m1), granzyme B (Mm00442837_m1), perforin (Mm00812512_m1) and IFNγ (Mm01168134_m1). All genes were normalized against GAPDH (Mm99999915_g1).

### Granzyme B protein analysis in lung samples

Right lung caudal lobes were homogenized by TissueLyser LT and processed as per the manufacturer’s protocol (Qiagen). Granzyme B protein was quantified by ELISA according to the manufacturer’s protocol (eBioscience).

### *In vitro* stimulation of bone marrow monocytes

Bone marrow cells were harvested from both femurs and tibias. Monocytes were isolated using an EasySep™ Mouse Monocytes Isolation Kit (STEMCELL, Vancouver, BC, Canada). 30000 monocytes plated per well in a round bottom 96 well plate in RPMI 1640 with 10% FPS and penicillin-streptomycin. Cells were either unstimulated or challenged with LPS (1ug/mL; LPS from E. coli O111:B4; Sigma-Aldrich) for 24 hours. Supernatant was collected and IL-1β levels were measured by ELISA (R&D systems, Minneapolis, MN, USA).

### Infection models

*Pseudomonas aeruginosa* (PA14), *Streptococcus pneumoniae* (P1121) and *Salmonella enterica (typhimurium)* (CDC 6516-60) were individually grown aerobically overnight in Luria Broth (LB), pelleted, and resuspended in PBS to the appropriate concentration based on optical density (OD600). Mice were treated with one of QBKPN, QBECO, QBSAU or vehicle starting 14 days before bacterial challenge and administrations were given every second day throughout the experiment, except for the day of bacterial challenge. On day zero, mice were challenged with the appropriate bacteria. For the lung infection models, mice were challenged with *S. pneumoniae* or *P. aeruginosa* by intranasal instillation of 5.0×10^5^ CFU[38]. For the intraperitoneal injection models, mice were challenged with 1 × 10^6^ CFU of *S. enterica (typhimurium)* bacteria by IP injection [39,40]. For the skin infection model, mice were challenged with *P*. *aeruginosa* by intradermal injection of 6.5×10^6^ CFU of bacteria [41]. On day three post-challenge, mice were sacrificed and the targeted organ (skin, lungs, or spleen) was aseptically resected, weighed, homogenized, and assessed for bacterial load. Counts were performed on LB agar plates for *S. pneumoniae, Hektoen* enteric agar plates[42] (Becton Dickinson, Franklin Lakes, NJ, USA) for *S. enterica (typhimurium)* and *Pseudomonas* selection agar plates (Teknova, Hollister, CA, USA) for *P. aeruginosa*.

### Data Analysis

GraphPad Prism 6 Software (GraphPad Software, San Diego, CA, USA) was used to perform statistical analyses. Data are expressed as mean ± SD, except for serum cytokines levels, which are expressed as mean ± SEM. For comparison between 2 groups, Student’s *t*-test was performed. For comparison between multiple groups, one-way ANOVA analysis followed by a post-hoc multiple comparisons test was conducted using a Holm-Sidak’s test. For comparison between multiple groups at multiple time points, a two-way ANOVA analysis followed by Tukey’s multiple comparison test was performed. Differences in survival time was determined by Log-rank (Mantel-Cox) analysis. Outliers in tissues cytokines were identified by GraphPad’s ROUT test. Statistical significance was set at α level of 0.05.

## ACKNOWLEDGEMENTS

We thank Dr. Brent Berwin and Dr. George O’Toole at Dartmouth College for their kind gifts of *P. aeruginosa* PA14, *S. pneumoniae* P1121 and *S. enterica (typhimurium)* CDC 6516-60. We thank Dr. Jeff Schlom at NCI for his kind gift of MC-38 cells, and Dr. Dietrich Kabelitz at the University Hospital Schleswig-Holstien for his kind gift of Panc02 cells. We thank Rebecca Anderson and Karl Lau for their technical support. We also thank Dr. Bruce Vallance for his critical review of the manuscript and for excellent scientific discussions.

## CONTRIBUTIONS

S.K. and M. Bazett designed the experiments, interpreted the data and wrote the first draft of the manuscript. H.P.S., M. Bosiljcic, B.L., S.D., S.W.Y.W. conducted the experiments. M. Bosiljcic also contributed to designing and optimizing the cancer models and H.P.S. helped design and optimize the DSS/AOM cancer model and *in vitro* experiments. A.M.C. and D.W.M. performed the infection model experiments. M.G.N. helped design the experiments showing trained innate immunity. H.G. conceived the idea and developed the microbial-based treatments described. All authors reviewed and edited the final version of the manuscript.

## Supplemental Figure Legends

**Supplemental Figure 1: Anti-cancer efficacy is organ specific, but not cancer type specific. (A)** Lung surface tumor count in mice 17 days after seeding with B16F10 melanoma cells via tail vein injection, and administered Vehicle, QBKPN and QBECO every second day by subcutaneous injection, starting 10 days before tumor seeding. n = 9-10 mice per group. Mean ± SD shown. ** p < 0.01 (one-way ANOVA with Tukey’s multiple comparisons). **(B)** Survival curve of mice seeded by interperitoneally injection with Panc02 pancreatic ductal adenocarcinoma cells and administered Vehicle, QBKPN and QBECO every second day by subcutaneous injection, starting 10 days before tumor seeding. n = 10 mice per group. *** p < 0.005, Log-rank test.

**Supplemental Figure 2: Distribution of fluorescently labelled QBKPN after 3 injection. A** Fluorescent intensity maps of *in vivo* optical image of Cy5.5-QBKPN after injections 1, 2 or 3 injection. Injection were given at 0 hours, 48 hours and 96 hours. n = 3 mice. **(B)** Pharmackinetic data depicting the presence of Cy5.5-labeled QBKPN (cells/mL) in blood after a single Cy5.5-QBKPN injection. Blood was collected from three mice at rotating timepoint with each point represent the value from a single mouse.

**Supplemental Figure 3: Cytokine changes in the lung tissue in cancer bearing mice at day 5 and day 17.** Cytokines were measured in lung homogenates in the B16F10 lung cancer model at 5 and 17 days post tumor inoculation. Cytokines were measured by 31-cytoplex. GM-CSF (day 5), IL-1α, IL-1β, Il-4, IL-9, IL-12p40, IL-17, CXCL1, M-CSF, MIP1α, MIP1β (day 5) and TNFα were detected but no significant changes were measured. GM-CSF (day 17), Il-3, IL-5, IL12p70 and MIP-1β (day 17) were measured but were not detectable. Data represents mean ± SD. n=8-10 mice per group. * p < 0.05; ** p < 0.01; *** p < 0.001; **** p < 0.0001, two-way ANOVA, Tukey’s multiple comparison test.

**Supplemental Figure 4: Cytokine changes in the lung tissue in cancer free mice after 27 days of treatment.** Cytokines were measured in lung homogenates in cancer free mice at 27 days after start of vehicle, QBKPN or QBECO treatment. Cytokines were measured by 31-cytoplex. GM-CSF (day 5), IFNγ, IL-1β, IL-6, IL-9, IL-7, IL-10, IL-15, MIP1α, MIP1β and VEGF were detected but no significant changes were identified. Il-3, IL-5 and IL12p70 and MIP-1β measurement were not detectable. Data represents mean ± SD. n=8-10 mice per group. * p < 0.05; ** p < 0.01; *** p < 0.001; **** p < 0.0001, one-way ANOVA with Holm-Sidak’s post-hoc test.

**Supplemental Figure 5: Cytokine changes in the serum after a single infection of vehicle, QBKPN and QBECO.** Cytokines were measured in serum in cancer naïve mice 5, 12 and 24 hours after a single subcutaneous injection of vehicle, QBKPN or QBECO. Cytokines were measured by 31-cytoplex. Eotaxin-1, IL-2, Il-1α, IL-7, IL-9, and VEGF were detected but no significant changes were identified. IFNγ, IL-3, IL-4, and LIF measurement were not detectable. Data represents mean ± SD. n=8-10 mice per group. * p < 0.05; ** p < 0.01; *** p < 0.001; **** p < 0.0001, one-way ANOVA with Holm-Sidak’s post-hoc test. Data from 5 hours in vehicle and QBKPN treated mice has been previously published[8].

**Supplemental Figure 6: QBSAU has anti-cancer efficacy in skin cancer models, but not in the lung and intraperitoneal cancer models. (A)** Skin tumor size in a mouse model where B16F10 cells are seeded dorsally via subcutaneous injection. Vehicle, QBKPN, QBECO and QBSAU administered by ventral subcutaneous injected every second day starting 10 days before tumor seeding and continued throughout the experiment. Mean ± SD; n=5 per group. ** p < 0.01, one-way ANOVA with Holm-Sidak’s post-hoc test. **(B)** Lung tumor count in the RFP-LLC mouse lung cancer model at day 15 mice administered with vehicle and QBSAU. Vehicle data reproduced from Fig. 1A. **(C)** Survival curve in the MC-38 mouse model in mice administered with vehicle and QBSAU. Vehicle data reproduced from Fig. 1B.

**Supplemental Figure 7: Demonstration that organ-specific pathogen protection is independent of disease-causing pathogen.** *P. aeruginosa* count in the lungs of mice 3 days after bacterial challenge by intratracheal instillation. Mice were treated with Vehicle, QBKPN or QBECO every second day starting 14 days before bacterial challenge. n = 8 mice per group. Mean ± SD shown. ** p < 0.01; **** p < 0.0001, one-way ANOVA, with Holm-Sidak’s post-hoc test.

